# Decoding stress-induced adaptation of cancer cells in spatial confinement by integrating live-cell imaging and mathematical modeling

**DOI:** 10.64898/2026.11.01.763387

**Authors:** Gopal Vyankatesh Kulkarni, Atish Mohanty, Ayalur Raghu Subbalakshmi, Arin Nam, Sourabh Kotnala, Dana Do, Prakash Kulkarni, Mohit K. Jolly, Supriyo Bhattacharya, Ravi Salgia

## Abstract

The adaptive strategies employed by cancer cells to combat stress when grown in 2D cultures have been extensively studied, including applying evolutionary game theory. However, relatively little is known about these strategies when the cells are grown in confinement that resemble the tumor microenvironment more closely. Here, we used cisplatin-sensitive and -tolerant lung cancer cells that were grown as monotypic or heterotypic spheroids in a microfluidics system and monitored in real time. Competition and cooperation between these cells were modeled by estimating the parameters through the best fit of observed measurements using a modified version of the Cellular Potts model. The results indicated that the two sensitive and tolerant populations exhibited different cooperative and competitive strategies under changing environmental conditions. Importantly, we found that in addition to initial cell type composition, the population fitness was governed by the spatial organization, intercellular interactions, and dynamic remodeling of the local microenvironment by these cells. Further, tolerant cells exhibited similarities to a persister trait and appeared to cooperate with sensitive cells, enabling them to evade chemotherapy.

## Introduction

Cancer is a remarkably adaptive disease, driven by its ability to evade environmental challenges and enhance cellular fitness under adverse conditions(*1, 2*). This adaptability reflects the evolutionary legacy of cells, which retain the cumulative “experiments” of billions of years of evolution. These experiences manifest as “latent attractors,” enabling cancer cells to transition between phenotypic states in response to stress or environmental changes(*3*).

Although cancer has traditionally been viewed as a genetic disease, growing evidence underscores the critical role of non-genetic mechanisms, such as epigenetic reprogramming, metabolic plasticity, and stochastic fluctuations, in driving cancer progression and therapeutic resistance(*4-8*). Together, genetic and non-genetic mechanisms create a duality that mirrors evolutionary processes and contributes to the emergence of drug resistance—one of the greatest challenges in oncology(*9, 10*).

Drug resistance arises through diverse pathways, combining genetic mutations and non-genetic adaptations that enable cancer cells to survive pharmacological stress. In vitro cancer cell lines have been instrumental in unraveling these mechanisms, serving as models for both single-cell responses and collective behaviors such as cooperative survival and migration(*6-8, 11-13*).

Despite the utility of 2D culture systems in drug resistance studies, these systems fail to replicate the complexity of the tumor microenvironment (TME), which profoundly influences cancer behavior. The TME is characterized by gradients of oxygen, nutrients, and metabolites, as well as interactions with stromal cells such as fibroblasts and immune cells(*14, 15*). 3D culture systems, including spheroid and organoid models, have emerged as essential tools for studying cancer in more physiologically relevant conditions. These systems help recreate key TME features, like hypoxia, nutrient diffusion gradients, and extracellular matrix stiffness, providing a more accurate representation of in vivo tumors(*16-18*).

In our previous work, using cisplatin-sensitive and tolerant non-small cell lung cancer (NSCLC) cells cultured in 2D, we demonstrated the role of non-genetic mechanisms in drug tolerance(*19*). The dynamics of drug-sensitive and drug-tolerant cells in homogeneous 2D systems has been extensively studied(*20, 21*); however, which strategies the cell populations follow in spatial confinement remains elusive(*22, 23*). Thus, to investigate this group dynamics in a more realistic setting, we transitioned to 3D spheroid systems using ultra-low attachment plates designed to promote spheroid formation. Our experimental setup was motivated by a microfluidics device (death galaxy) designed by Austin and co-workers to create spatially confined environments mimicing the TME(*24*). The growth and spatial configurations of sensitive and tolerant cells in the microfluidic apparatus were monitored in real time using the IncuCyte® Live Cell Analysis System.

To complement these experiments, we employed CompuCell3D (CC3D), a computational platform for simulating complex biological systems(*25*). CC3D integrates diverse mathematical models, including the cellular Potts model, to simulate cell clustering, growth, division, death, and chemotaxis. Additionally, it enables modeling of reaction-diffusion dynamics, such as nutrient gradients, and automata rules for cellular differentiation. This integrated experimental-computational approach provided novel insights into the phenotypic plasticity of cancer cells, revealing how non-genetic mechanisms drive group behavior and promote the emergence of drug tolerance.

## Results

### Cocultured sensitive and tolerant cells dynamically alter their spatial organization in response to changing population

The confined environment of the TME is a critical factor in shaping tumor ecology, driven by the adaptability and heterogeneity of cancer cells and their spatial interactions. To determine whether the spatial behavior of the sensitive and tolerant cells differ in a confined environment, we co-cultured them starting with different seeding ratios (and a constant starting population of 5000 cells), monitored over a period of 6 days. For comparison, mono-cultures of these cell types were also monitored over the same period (Fig. 1A, schematic and representative images obtained using IncuCyte live cell imaging). Under mono-culture condition, the sensitive and the tolerant cells exhibited distinct spatial behavior with sensitive cells forming a diffused cluster occupying the entire 3D space, and the tolerant cells forming a tight spheroid despite having room for expansion. Moreover, the fluorescence intensity from the sensitive cells progressive decreased over a period of 6 days. In contrast, the fluorescent intensity from the tolerant cell remained sustained throughout the coculture period. These observations were supported by quantitative measurement of fluorescent intensity over time that served as a proxy for cellular health and population (Fig. 1B). Based on these observations, it can be concluded that the sensitive cell population declined, while the tolerant population increased over time.

**Figure 1.**
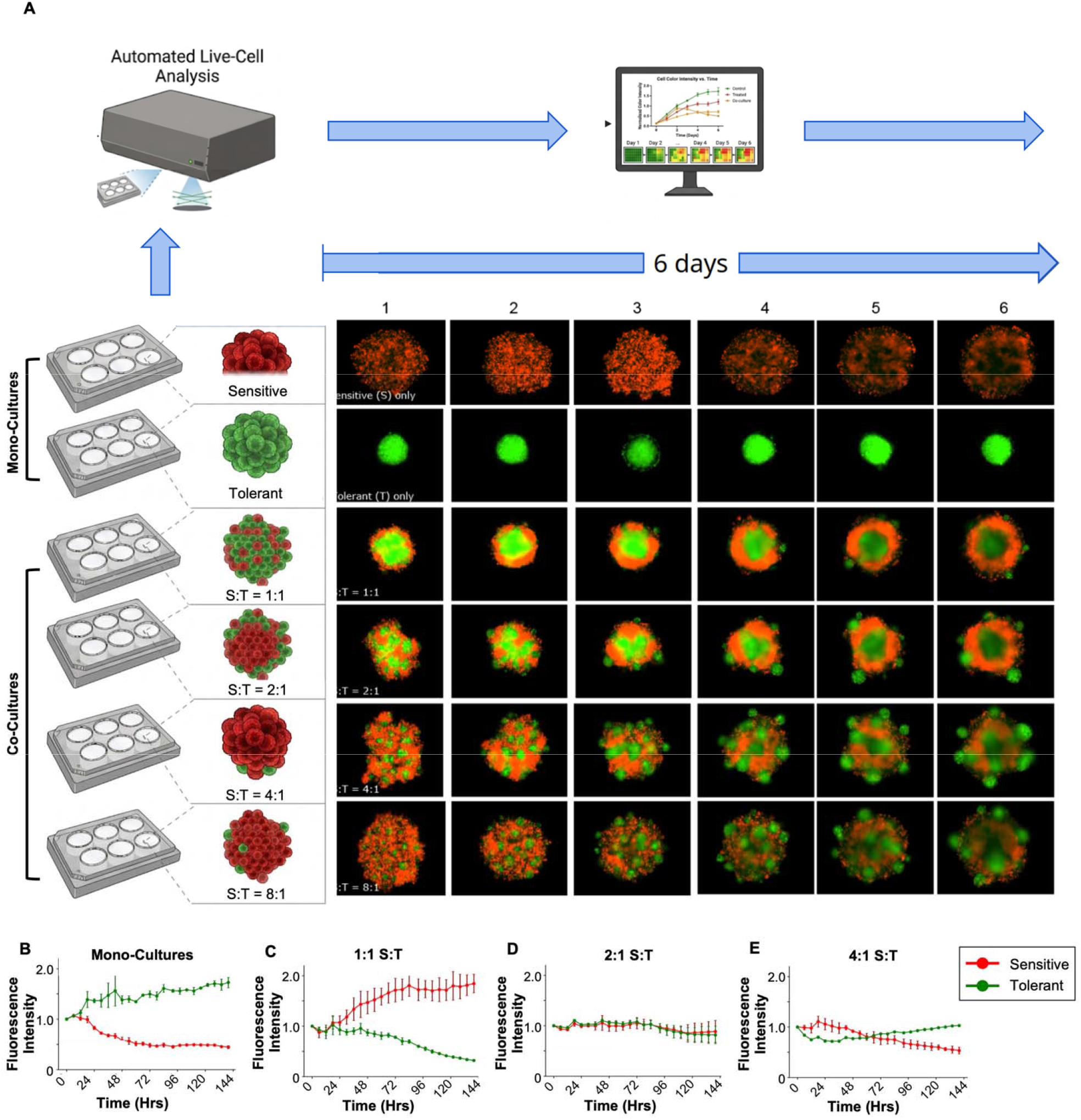
Real-time monitoring of sensitive and tolerant cancer cell dynamics *in vitro*. **(A)** Schematic illustration of the experimental setup utilized for the real-time fluorescence monitoring of sensitive and tolerant cancer cell cultures, along with live cell imaging showing temporal evolution of fluorescence intensity in monocultures and co-cultures. **(B–E)** Quantitative florescence intensity over time measured by IncuCyte Live cell monitoring system, depicting growth dynamics across varying initial seeding ratios of sensitive to tolerant populations, specifically 1:1, 2:1, and 4:1. Data are presented as mean ± standard deviation (n = 3 independent experiments).

We next compared the mono-culture observations with co-culture experiments seeded with different ratios of sensitive and tolerant cells, while keeping the total cell number constant (5000 cells/ well). When co-cultured, the sensitive and tolerant cells showed diverse spatial organizations that varied both temporally and with changing seeding ratios (Fig. 1A). At a 1:1 (sensitive:tolerant) seeding ratio, the tolerant cells formed a tight spheroid similar to the mono-culture. However, the sensitive cells, instead of dispersing, clustered around the tolerant cells, suggesting that the presence of tolerant cells had altered their spatial behavior. Over time, the layer of sensitive cells increased in thickness, while the fluorescent intensity from the tolerant cells decreased, suggesting declining population. The temporal fluorescence measurements confirmed this, showing progressive increase and decrease over time for the sensitive and tolerant cell populations respectively (reversed trends from monoculture, Fig. 1B).

Increasing the seeding ratio of sensitive: tolerant to 2:1 led to a new spatial organization on day 1, where the sensitive cells still surrounded the tolerant cells, but instead of forming a homogenous layer, these cells surrounded micro-clusters of tolerant cells on the surface of the tolerant spheroid. Then, starting day 3, the sensitive cells merged into a single layer that increased in thickness over time. The tolerant cells remained mainly confined within a tight spheroid, except 3-4 micro-clusters that emerged outside the sensitive cell layer. Both cell populations remained stable over time, as indicated in Fig. 1B. Increasing the seeding ratio to 4:1 and 8:1 led to sensitive cells entirely covering the surface of the tolerant spheroid on day 1 and micro-clusters of tolerant cells appearing on day 3. Over time, the fluorescent intensity from the sensitive cells decreased, suggesting declining cell population (and supported by temporal fluorescence measurements, Fig. 1B). While the central core population of tolerant cells surrounded by the sensitive cells decreased in intensity, the micro-clusters outside the sensitive cell layer showed increased intensity, resulting in an overall increase in tolerant cell population with time (Fig. 1B). In summary, the confined microenvironment and cell interactions strongly influenced how the sensitive and tolerant cells organized spatially and survived.

### Multiscale agent-based simulations recapitulate the group dynamics of sensitive and tolerant cell co-cultures

To understand the ecological principles governing this experimentally observed collective behavior, we developed a computational model of cellular ecosystem in confinement using a multiscale agent-based computational framework based on the Cellular Potts model (further elaborated in the Methods section). In this model, space is discretized into a lattice grid, where individual cells occupy localized regions of adjacent pixels and the empty space constitutes the microenvironment, also referred to as the medium (Fig. 2A). The model incorporates two critical aspects of the observed cellular behavior – a) the spatial clustering and b) variable growth dynamics. The spatial clustering is governed by an interaction energy term that determines how strongly the two different cell types (sensitive and tolerant) cluster together (table 1). For example, tolerant-tolerant and sensitive-tolerant interaction energy is lower (more favorable) compared to sensitive-sensitive interaction based on the observed clustering behavior in the mono- and co-culture experiments (Fig. 1A). Cellular growth dynamics is modulated by the changing microenvironment condition over time. Given our experimental observations of decrease in sensitive cell populations in the mono-culture experiments, we hypothesized that proliferating sensitive cells secrete diffusible factors in the microenvironment that are toxic to the cell population. These could be metabolic byproducts or other cytotoxic factors that we collectively refer to as “stress”(*26, 27*). Conversely, our experiments show that the tolerant cell population under monoculture conditions increased over time, potentially implying that diffusible factors secreted by these cells favor cellular growth. We refer to these secreted molecules that can be the basis for cooperative behavior by tolerant cells, as “co-op”. The numerical values of stress and co-op related parameters were optimized based on agreement with observed spatial and growth trends and are provided in table 2.

**Table 1.** Pairwise contact energy parameters.

| **Pair** | **Contact Energy (J)** |
|---|---|
| Medium–Medium | 10.0 |
| Medium–Sensitive | 10.0 |
| Medium–Tolerant | 10.0 |
| Sensitive–Sensitive | 16.0 |
| Sensitive–Tolerant | 18.0 |
| Tolerant–Tolerant | 15.0 |

**Table 2.** Parameters governing the secretion and diffusion of stress and co-op.

| **Parameter** | **Stress** | **Co-op** |
|---|---|---|
| Global diffusion constant | 0.0001 | 0.001 |
| Global decay constant | 0.0001 | 0.00001 |
| Sensitive specific diffusion | 0.1 | 0.1 |
| Tolerant specific diffusion | 0.1 | 0.1 |
| Sensitive specific decay | 0.0001 | 0.0001 |
| Tolerant specific decay | 0.0001 | 0.0001 |
| Secretion by | Sensitive = 10.0 | Tolerant = 10.0 |
| Boundary conditions | homogeneous Neumann | homogeneous Neumann |

**Figure 2.**
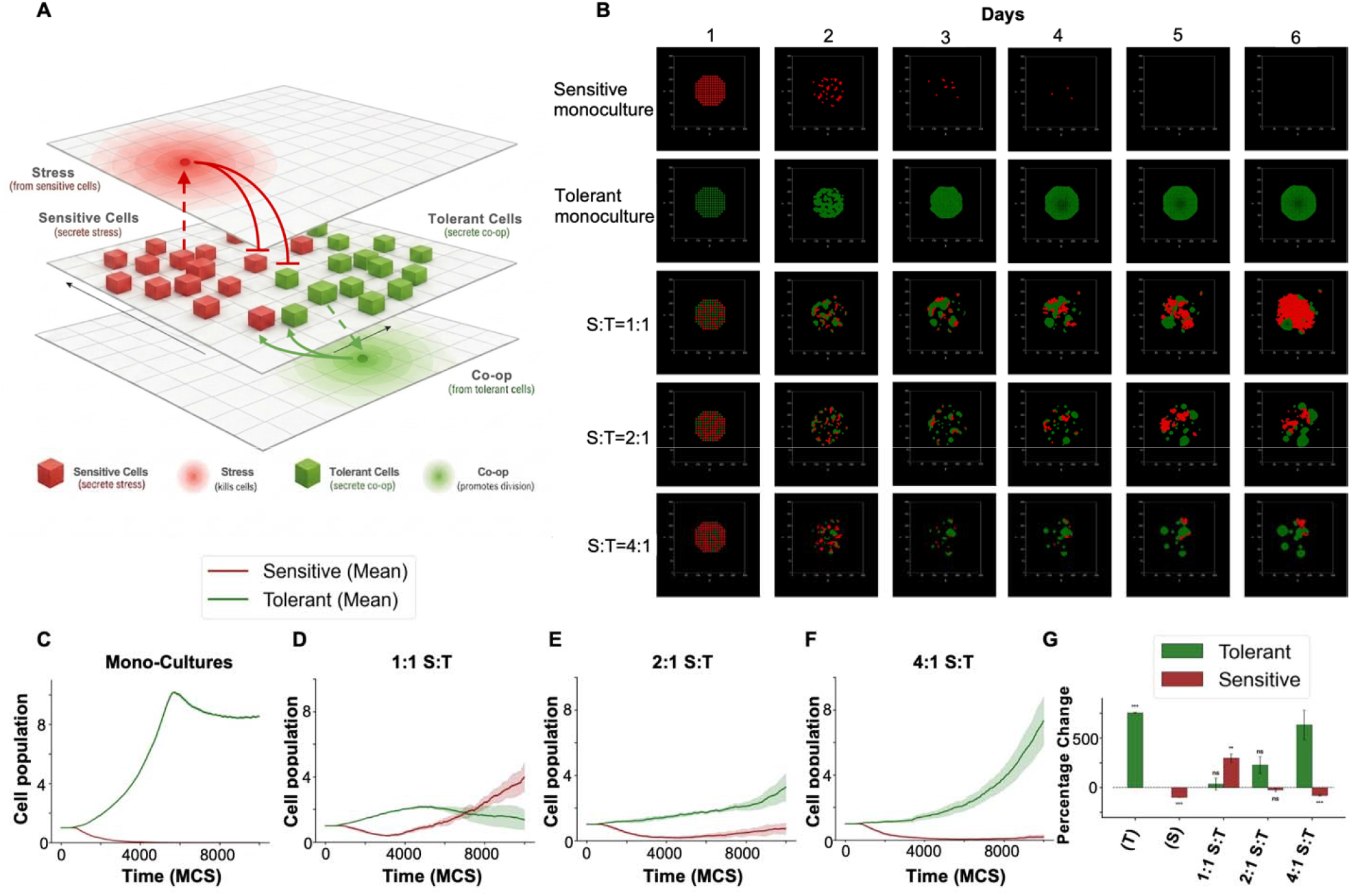
Multiscale agent-based modeling (ABM) of tumor cell population dynamics. **(A)** Schematic overview of the multiscale agent-based model architecture detailing the computational framework. **(B)** Representative ABM simulation snapshots of sensitive and tolerant cell distributions in corresponding monocultures and co-cultures at 10,000 Monte Carlo Steps (MCS). **(C–F)** Average population dynamics of specific cell types tracked over the duration of the simulation (n = 5 independent simulation runs). **(G)** Percentage change in cell population relative to the initial baseline for each respective cell type across the simulated culture conditions.

At any given point, the rate of cell population growth depends on the joint level of stress and co-op in the medium. Both cell types encounter rapid cell death under high stress (> 5000 units) and proliferate under high co-op beyond pre-defined thresholds (> 10,000 units for sensitives and > 7000 units for the resilient tolerant cells). Additionally, given the high fitness of tolerant cells in co-culture with the high sensitive cell population (e.g. at 4:1 S:T ratio), we postulated that these cells would be able to proliferate under elevated stress, if the co-op level is also high. The rules for proliferation and cell death under different stress and co-op regimes are described in tables 3 and 4 (also see methods).

**Table 3.** Stress and co-op dependent rules for division and death of sensitive cells.

| **Event** | **Condition** | **Per-MCS probability** |
|---|---|---|
| Division | co-op > 10,000 | 0.003 |
| Death | stress > 5000 | 0.001 |

**Table 4.** Stress and co-op dependent rules for division and death of tolerant cells.

| **Event** | **Condition** | **Per-MCS probability** |
|---|---|---|
| Division (stress-independent) | co-op > 7000 | 0.0003 |
| Division (low-stress refuge) | stress < 2000 and co-op > 2000 | 0.0003 |
| Death | stress > 5000 | 0.001 |

In CompuCell3D simulations, each system was initialized with a fixed population of sensitive and tolerant cells at different seeding ratios and simulated for 10,000 steps (∼ 6 days) using the monte carlo (MC) Metropolis algorithm. The simulation at each seeding ratio was replicated five times with a different random seed to obtain the average growth trends and error margins. The simulation snapshots captured at regular intervals describe the temporal evolution of spatial configurations adopted by both cell-types (Fig. 2B). In the monoculture condition, sensitive cells remained dispersed with decreasing population over time, while the tolerant cells formed a tight cluster with increasing population, as observed in the experiments. Under co-culture conditions at different seeding ratios (8:1 ratio was not simulated due to spatiotemporal similarity with 4:1), the sensitive cells surrounded the tolerant cells, which maintained their clustering tendency, while forming smaller individual clusters. The propensity to form smaller homotypic clusters by tolerant cells increased at higher sensitive: tolerant (S:T) seeding ratios (Fig. 1A). This trend is in line with the experimental behavior, where higher S: T seeding ratios of 2:1 and 4:1 exhibited increased tendency of tolerant cells to form micro-clusters (Fig. 1A).

The spatial observations are also qualitatively in agreement with the experimental results with some notable differences. For example, at 1:1 seeding ratio, the tolerant cells formed more homotypic clusters compared to experiments. Also, at 1:1 and 2:1 S:T ratios, there were not enough sensitive cells to form a circular ring around the tolerant cells, as observed in experiments. We envision that closer agreement with the experimental spatial configurations would require simulating significantly larger cellular populations that is beyond the computational capacity of *in-silico* model. However, the temporal growth trends derived from the simulations (Fig. 2C-F) agreed with the experimental ones (Fig. 1B-E. The percent change in the cell population at 6 days, calculated from the replicate simulations, reproduced the striking reversal in the population trends of the two cell types observed in experiments, e.g., from monoculture (sensitive population frequency: low, tolerant population frequency: high) to 1:1 coculture condition (sensitive population frequency: high, tolerant frequency: low; Fig. 2G). However, increasing the sensitive cell proportion to 4:1 caused the tolerant cells to regain a higher dominance in the population. To explain the basis of this dynamic change in behavior of the two cell types with seeding ratio, we next analyze in our simulations the cell interactions and microenvironment changes over time, as described in the next section.

### Microenvironment changes and spatial organization govern fitness dynamics in sensitive-tolerant cocultures

To elucidate the ecological interactions between sensitive and tolerant cells, we examined two key features of our model system: cell-cell interactions and the local concentrations of diffusible factors (stress and co-op) and assessed how these features evolved over time and across different S:T seeding ratios. A distinguishing feature of the coculture system was the occurrence of heterotypic interactions between sensitive and tolerant cells, in addition to monotypic interactions within each cell type. The fraction of heterotypic contacts, defined as instances in which a sensitive and a tolerant cell occupied adjacent lattice sites, relative to monotypic contacts decreased with increasing S:T ratio (Fig. 3A). Together with the reduced fitness of sensitive cells at higher S:T ratios, this relationship suggests that heterotypic contact with tolerant cells favors sensitive-cell fitness, whereas increased monotypic interactions among tolerant cells are associated with greater tolerant-cell fitness (Fig. 3A).

**Figure. 3.**
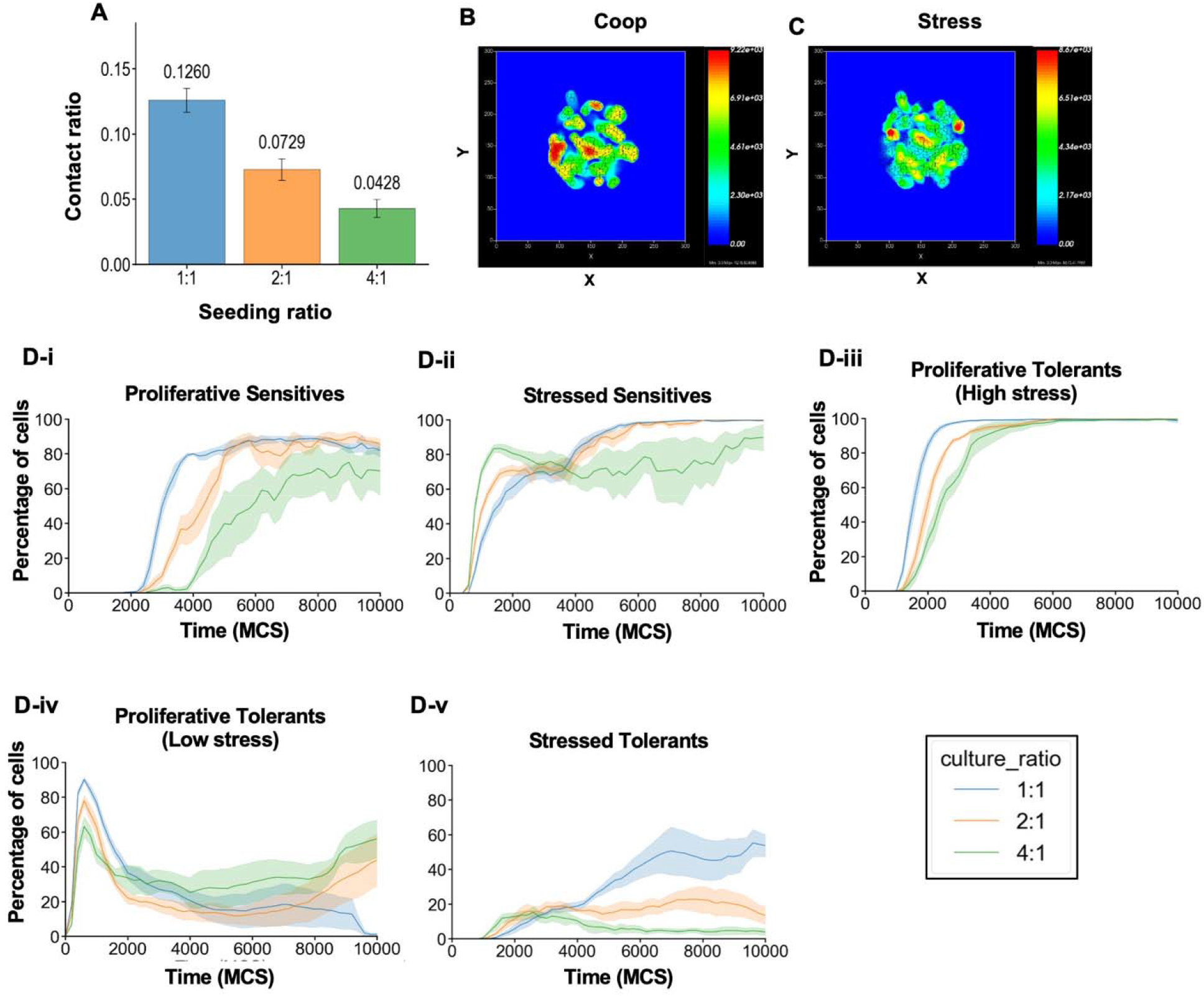
Microenvironmental field transitions in simulated co-cultures. (A) Ratio of heterotypic to homotypic cell–cell contacts across varying initial seeding ratios (1:1, 2:1, and 4:1). (B, C) Representative spatial visualizations of the microenvironmental fields driving cellular behavior, displaying the (B) cooperative (co-op) field and (C) stress field. (D) Temporal evolution of distinct phenotypic subpopulations across the 1:1, 2:1, and 4:1 seeding ratios (n = 5 runs per condition). Panels show the percentage of the (i) proliferative sensitive population, (ii) stressed sensitive population, (iii) proliferative tolerant population, (iv) stressed tolerant population, and (v) proliferative tolerant population specifically residing in low-stress microenvironments.

We hypothesized that the beneficial effect of heterotypic interactions on sensitive cells arises, at least in part, from co-op secreted by tolerant cells. In our simulations, cell proliferation was governed by the competing effects of co-op and stress, generating distinct proliferative niches for the two cell types (Fig. S1A-B). Briefly, both cell types proliferated under high co-op conditions and experienced increased cell death under high stress. However, tolerant cells could proliferate at higher stress levels and lower co-op concentrations than sensitive cells, resulting in distinct low- and high-stress proliferative regimes (Fig. S1B).

To examine how the simulated populations traversed this ecological landscape, we first characterized the spatial distributions of co-op and stress over time (Fig. 3B-C) and then projected individual simulation timepoints onto the joint stress/co-op landscape (Fig. S1C). Comparison of these trajectories across seeding ratios revealed composition-dependent differences in population dynamics. At a 1:1 S:T ratio, for example, the system frequently occupied regions characterized by both high stress and high co-op, corresponding to conditions that supported high proliferation but were accompanied by substantial cell death. In contrast, at a 4:1 ratio, the system remained predominantly within the favorable low-stress region (gray area), with only occasional excursions into the high-stress regime.

To further resolve these fitness dynamics, we quantified the fraction of each cell population occupying the proliferative and high-stress regimes over time (Fig. 3D). Consistent with the high proliferation observed at the 1:1 seeding ratio, more than 80% of the sensitive-cell population rapidly entered the proliferative regime and remained there for the remainder of the simulations (Fig. 3D-i). By contrast, at the 2:1 and 4:1 ratios, sensitive cells exhibited delayed entry into the proliferative regime (Fig. 3D-i) and more rapid entry into the high-stress regime (Fig. 3D-ii), consistent with their reduced fitness under these conditions.

Tolerant cells exhibited markedly different dynamics. Across seeding ratios, a large fraction of the tolerant population occupied the proliferative regime, whereas a smaller fraction entered the high-stress regime (Fig. 3D-iii, iv). Notably, at the 1:1 ratio, an increased fraction of tolerant cells entered the high-stress regime (Fig. 3D-v). This increase coincided with stress generated by the rapidly proliferating sensitive-cell population and provides a mechanistic explanation for the reduced fitness of tolerant cells under this condition. Conversely, at the 2:1 and 4:1 ratios, fewer tolerant cells occupied the high-stress regime, consistent with their increased proliferation.

The observation that tolerant cells experienced lower stress at higher S:T ratios, despite the larger initial proportion of stress-producing sensitive cells, can be understood by considering the spatial organization of the populations (Figs. 1A and 2B). In monoculture and 1:1 coculture, tolerant cells formed a single compact cluster that was surrounded by sensitive cells. This spatial arrangement restricted tolerant-cell proliferation while simultaneously benefiting sensitive cells by creating a localized region of high co-op concentration. In contrast, at higher S:T ratios, particularly 4:1, tolerant cells formed multiple smaller clusters. This dispersed organization reduced the access of sensitive cells to regions of concentrated co-op, thereby limiting their proliferation and promoting their loss from the population. The resulting decline in the sensitive-cell population, together with the ability of tolerant cells to proliferate under relatively high-stress conditions, ultimately favored expansion of the tolerant population.

Collectively, these results indicate that the population fitness is determined not simply by the initial abundance of sensitive and tolerant cells, but by an interplay among spatial organization, heterotypic and monotypic interactions, and dynamic remodeling of the local stress/co-op microenvironment. By capturing these coupled spatial and ecological processes, our multiscale model provides a mechanistic framework for explaining the otherwise counterintuitive population behaviors observed in the death galaxy experiments.

### Co-culture with tolerant cells improved sensitive cell survival in presence of the chemotherapeutic drug cisplatin

We next examined how cisplatin treatment perturbs the population balance between sensitive and tolerant cells under co-culture conditions. We compared the growth dynamics of sensitive and tolerant cells in monoculture and in 1:1 coculture, in the presence or absence of a fixed dose of cisplatin (2.5 μM). This concentration was lethal to sensitive cells but sublethal to tolerant cells (Fig. 4A–B).

**Figure 4.**
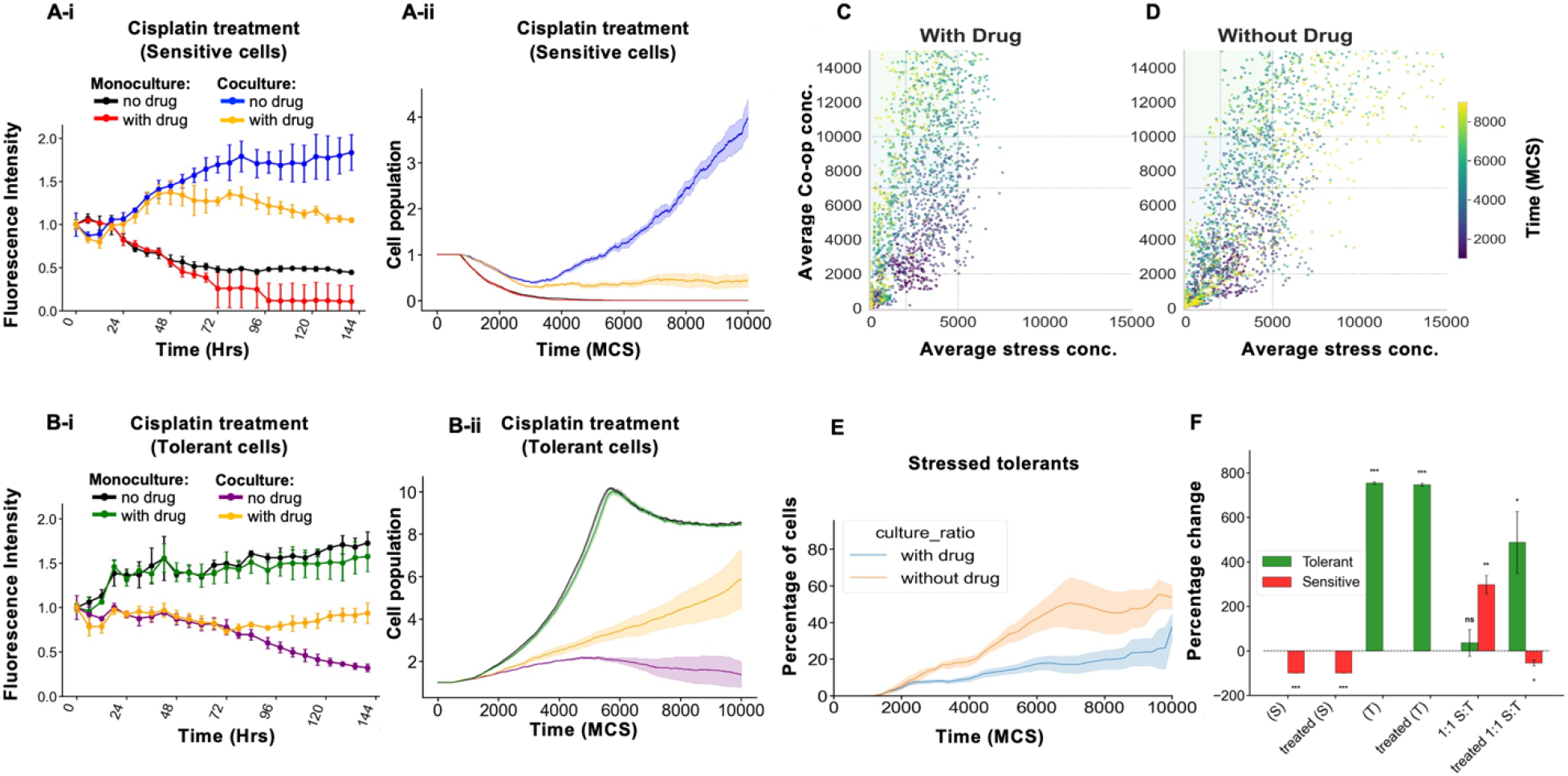
Experimental and simulated therapeutic responses of mono- and co-cultures to cisplatin treatment. (A) Treatment response of the sensitive cell population: (i) Fluorescence intensity of untreated and cisplatin-treated sensitive cells in experimental monocultures and 1:1 co-cultures; (ii) Corresponding ABM-simulated average cell populations for untreated and cisplatin-treated sensitive cells. (B) Treatment response of the tolerant cell population: (i) Fluorescence intensity and (ii) simulated average cell populations for untreated and cisplatin-treated tolerant cells in mono- and 1:1 co-cultures. (C, D) Scatter plots of simulation time-points as function of average stress and co-op concentrations in 1:1 co-cultures for the (C) cisplatin-treated and (D) untreated conditions, colored by simulation time. The stress and co-op thresholds defining cell death and proliferation regimes are represented by grey straight lines. (E) Percentage of the stressed tolerant population tracked over the simulation duration, comparing cisplatin-treated and untreated scenarios. (F) Percentage change in the population relative to the initial baseline for each cell type under treated and untreated conditions.

As expected, cisplatin markedly reduced the growth of sensitive cells in mono-culture (Fig. 4A-i), whereas tolerant cells were only modestly affected (Fig. 4B-i). Strikingly, however, sensitive cells exhibited substantially improved survival under cisplatin treatment when cocultured with tolerant cells compared with sensitive-cell monoculture (Fig. 4A-i). Although the sensitive-cell population began to decline after approximately 72 h, its abundance at day 6 remained significantly higher than that observed in monoculture. In contrast, cocultured tolerant cells initially exhibited a gradual decline both in the presence and absence of cisplatin, followed by recovery after approximately 72 h, coincident with the decline in the sensitive-cell population. By day 6, the fluorescence intensities of the two populations were comparable. Together, these observations indicate that cisplatin induced a marked ecological shift, changing the coculture from a sensitive-cell-dominant state to a more balanced composition of sensitive and tolerant cells.

We next asked whether our *in silico* model could reproduce this drug-induced shift. Cisplatin treatment was modeled as a dose-dependent increase in cell death, with the magnitude of the effect determined by the tolerance threshold of each cell type (see Methods). To mirror the experimental conditions, we simulated a drug concentration that was sublethal to tolerant cells but lethal to sensitive cells. The simulated population dynamics qualitatively recapitulated the experimental trends (Fig. 4A-ii and 4B-ii).

Projection of the simulated trajectories onto the stress–co-op landscape provided insight into the ecological mechanisms underlying these dynamics. Under cisplatin treatment, the system predominantly occupied the favorable, low-stress region of the landscape (Fig. 4C), whereas, in the absence of drug, the population traversed regions characterized by high stress and high co-op (Fig. 4D). Consistent with this shift, the fraction of the tolerant population experiencing the high-stress regime decreased following cisplatin treatment (Fig. 4E). This reduction was associated with suppression of the sensitive-cell population, thereby alleviating the stress imposed on tolerant cells. The resulting persistence of the tolerant population, in turn, maintained cooperative support for sensitive cells, enabling a fraction of the sensitive population to survive despite direct drug-induced cytotoxicity. Thus, cisplatin fundamentally reorganized the ecological balance between the two populations: direct suppression of sensitive cells reduced the stress experienced by tolerant cells, while persistence of tolerant cells maintained sufficient cooperative support to permit partial survival of the drug-sensitive population (Fig. 4F).

## Discussion

Here, we have explored the complex dynamics between drug-sensitive and - tolerant cancer cell populations, using integrated experimental observations and computational modeling to understand the mechanisms underlying their interactions. Our findings demonstrate that the differences in their proliferative capacities, adhesion properties, and environmental adaptability influence spatial organization and competitive outcomes. Here, we combine real-time imaging with CompuCell3D simulations, to provide new insights into the dominance of sensitive cells under most conditions due to their replicative potential and the strategies employed by tolerant cells, like high cooperativity with itself and the medium, in constrained environments.

We observed that the sensitive cells consistently exhibited a higher proliferation rate, enabling them to achieve spatial dominance and outcompete tolerant cells when the two populations were initially present at a 1:1 ratio. Even when tolerant cells constituted a large fraction of the initial population, sensitive cells progressively expanded their clusters and constrained the growth of tolerant cells, ultimately dominating the available space and resources. This observation is consistent with previous studies where it has been seen that proliferative dominance often confers a competitive advantage, particularly in environments with abundant resources(*28-30*). The ability of sensitive cells to outgrow tolerant populations without any initial numerical advantage mirrors findings in collective cancer cell migration, where cell-cell cohesion and collective dynamics can strongly influence tumor expansion(*28, 31*). Together, these findings highlight the importance of cellular properties such as cell adhesion and proliferation in shaping competitive interactions within heterogeneous tumor cell populations.

In contrast, tolerant cells relied more strongly on adhesion to the extracellular matrix to maintain spatial presence. This survival-oriented phenotype allowed them to persist within mixed populations, particularly under conditions in which proliferative expansion was constrained. As the proportions of tolerant and sensitive cells approached parity, expansion of the tolerant population plateaued, whereas sensitive cells continued to exploit the available space and resources more effectively. This behavior is consistent with previous observations of stress-adapted cancer cell states, which can display enhanced persistence under adverse conditions while remaining less competitive in environments favoring rapidly proliferating phenotypes(*32, 33*). The simulations further revealed a distinct spatial organization, with sensitive cells forming dense outer layers and tolerant cells confined to interior regions, providing mechanistic insights into tumor architecture. This arrangement resulted from the interplay of differential adhesion and proliferation, where sensitive cells clustered at the periphery, creating a barrier that isolated tolerant cells to the core. Such spatial heterogeneity is reminiscent of spatial heterogeneity observed in solid tumors, where proliferative cells dominate oxygenated outer regions, while stress-adapted cells can persist in hypoxic cores(*33, 34*).

These findings underscore the importance of phenotypic diversity in tumor dynamics. Sensitive cells drive rapid tumor growth through proliferative dominance, whereas tolerant cells contribute to long-term heterogeneity by demonstrating survival resilience. This interplay highlights the role of environmental constraints, such as nutrient and oxygen gradients, in shaping competitive dynamics and tumor composition(*33, 35, 36*). Future research should incorporate additional microenvironmental factors, such as angiogenesis, nutrient diffusion gradients, and immune populations to better mimic *in vivo* conditions. Investigating how vascular normalization or targeted therapies impact sensitive and tolerant populations could provide strategies to overcome therapy resistance. Furthermore, integrating metabolic modeling into computational frameworks could enhance our understanding of resource allocation’s role in phenotypic competition and tumor evolution. Collectively, this study highlights the potential of combined experimental-computational approaches to advance our understanding of interactions in a cancer ecosystem constituting diverse subpopulations, and their impact on collective outcomes such as resistance to therapies.

## Materials and Methods

### Cell lines and reagents

Cell lines (H23, H1993) were obtained from American Type Culture Collection (ATCC) (Manassas, VA, USA) and cultured in RPMI 1640 medium (Corning) supplemented with 10% fetal bovine serum (FBS), L-glutamine (2 mM), penicillin/streptomycin (50 U/ml), sodium pyruvate (1 mM), and sodium bicarbonate (0.075%) at 37°C, 5% CO2. Cisplatin was provided by City of Hope National Medical Center clinics (Duarte, CA, USA). Puromycin was purchased from Thermo Fisher Scientific (Waltham, MA, USA). Suberoylanilide hydroxamic acid (SAHA) was purchased from Selleck Chemicals (Houston, TX, USA).

### Live cell imaging and analysis

The cisplatin sensitive H23 and cisplatin refractory H2009 cells were used for this study. These cells were engineered to express the NucLight Red or Green respectively as mentioned in the previous study (15). These cells were counted for viable cells and were seeded at a density of 5000 cells/ well of a 96 well ultra-low attachment plate procured form S-Bio (PrimeSurface® 3D culture: Ultra-low Attachment Plates: 96 well). In case of the coculture the H23 and H2009 cells were seeded at different number to achieve the ratios of 1:1, 2:1, 3:1, 4:1 or 8:1, while keeping in consideration that the total cell count / well remains at 5000 cells / well. The volume of medium used for each well was 100uL. For the cisplatin assay, the drug was diluted to a concentration of 2X fo the desired concentration in complete medium, and 100μl was added extra to each well on top of the culture media of 100μl. The drug was added 24h after he initial seeding and for control instead of cisplatin DMSO was used. Images were acquired in real time using the IncuCyte® Live Cell Imaging System (Essen Bioscience). Total red and green intensity was measured using IncuCyte image analysis software and the data was feed into graph pad prism for generating the figures.

### Multiscale modeling framework

The cell population dynamics were simulated using a two-dimensional Cellular Potts Model (CPM) implemented in CompuCell3D (version 4.6.0) software(*25*). The lattice measured 300 × 300 pixels, and each simulation was run for 10,000 Monte Carlo steps (MCS). In the CPM, each biological cell is represented as a domain of contiguous lattice sites sharing a common index, and cell properties such as volume, surface area and chemotactic motion are determined from stochastic, stochastic, energy-biased attempts to copy the index of a site into a neighboring site. Copy attempts are accepted according to the Metropolis criterion applied based on the effective Hamiltonian, *H* given below.

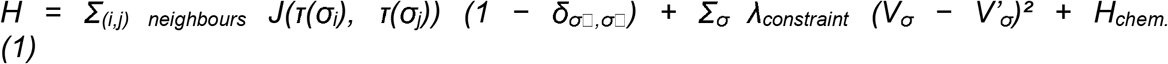

Where *J* is the contact (adhesion) energy between the cell types *τ* occupying neighboring sites, *V*_*σ*_and *V’*_*σ*_are the actual and target constraint of cell *σ* like volume or surface area, *λ*_*v*_is the constraint strength, and *H*_*chem*_ is the chemotaxis term.

### Agent types and properties

The model incorporates two distinct, heritable cell populations: sensitive (*S*, visualized in red) and tolerant (*T*, green). Each simulation is initialized with 145 cells placed on a regular, evenly spaced grid. Each cell type was initialized with a target volume of 50 lattice units. Initial population proportions and phenotypes were fixed at the start of the simulation, with phenotypic traits strictly inherited by all daughter cells. Simulations were initialized by seeding Sensitive and Tolerant cells into a spherical cluster (blob) of radius 70 lattice sites centered in the simulation box, utilizing a cell width of 7 units and an inter-cellular gap of 3 units **(Fig. S2)**. Contact energies for the cell type pairs are given in table 1.

### Field secretion

The stress field is secreted by sensitive cells and the coop is secreted by tolerant cells. These extracellular field concentrations were dynamically solved as reaction-diffusion PDEs of the form

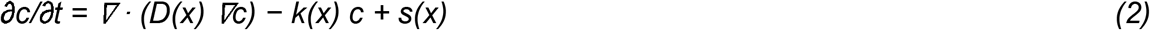

with the diffusion constant *D* and decay constant *k* taking different values in the medium and inside each cell type. The parameters are mentioned in table 2. Homogeneous Neumann (zero-flux) boundary conditions were applied on all four lattice edges.

### Proliferation and death

At MC step, the cell fate decision of division or death depends on the concentration of the diffusing fields sampled at the center of mass (COM) of each cell. Division or death takes place when the relevant field crosses the predetermined concentration threshold. Here Co-op concentration determines division of the cell and stress concentration decides the death of the cells.

A cell becomes mitotically competent only after completing the growth refractory period of 500 MCS since last division. Competent cells then divide stochastically with per-MCS probabilities for respective cell types, along a randomly oriented cleavage plane.

Proliferation incorporated two contrasting life-history strategies based on sensitivity to the co-op. Sensitive cells required a high co-op threshold to divide rapidly whereas tolerant cells required lower cooperative thresholds but divided at a reduced rate. Cellular mortality was governed by the local stress field. When stress levels exceeded the critical threshold both sensitive and tolerant cells experienced death at an identical stochastic rate (tables 3 and 4)

### Directed migration (chemotaxis)

Chemotaxis was applied to tolerant cells only. Tolerant cells were biased to move up the co-op gradient and down the stress gradient, implemented as an additional term in the change in effective energy for each copy attempt:

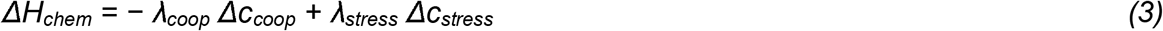

with chemotactic sensitivities *λ*coop and *λ*stress. Sensitive cells had no directed motility and were moved only by the CPM’s intrinsic membrane fluctuations.

### Drug treatment

Drug treatment was modeled as a global constant concentration *(C*_*drug*_ = 5.0) introduced after phenotype-specific burn-in periods (*MCS* > 2000 for sensitive cells and *MCS* > 1500 for tolerant cells). Drug-induced cytotoxicity was threshold-dependent and governed by a baseline kill rate of *k*_*kill*_= 0.00015*MCS*^-1^. Cells were susceptible to drug-induced death only when the drug concentration exceeded their respective resistance threshold (*sen_th* = 2.5 for sensitive cells; *tol_th* = 25.0 for tolerant cells). When this condition was satisfied, the per-MCS probability of cell death scaled linearly with the excess concentration (*k*_*kill*_ × [*C*_*drug*_ *-threshold*]), capped at a maximum of 1.0, and evaluated stochastically against a uniform random draw at each time step.

### Statistical Analysis

Percentage change values relative to initial conditions (*t* = 0) were calculated for both Tolerant and Sensitive populations across all mono-culture and co-culture configurations. To determine whether individual experimental conditions exhibited a statistically significant change from baseline, one-sample t-tests against a theoretical mean of zero were performed for each group using raw replicate data (*n* = 5 per configuration). Data are presented as means ± standard error of the mean (SEM). Statistical significance was evaluated at thresholds of * *p* < 0.05, ** *p* < 0.01, and *** *p* < 0.001, with nonsignificant results denoted as ns.

## Author contributions

Conceptualization: AM, PK, MKJ, SB and RS; Methodology: SB, AM, PK; Formal analysis: GVK, AM, ARS, PK; Resources: RS, MKJ; Data curation: AM, DD, AM; Writing – Original draft: PK, ARS, AM, SB; Writing – Review & Editing: GVK, AM, PK, MKJ, SB, RS; Supervision: MKJ, SB, RS; Project administration: PK, MKJ, RS; Funding acquisition: MKJ, RS.

## Conflict of Interest Statement

The authors declare no competing financial or non-financial conflicts of interest.

Mohit Kumar Jolly is Editor-in-Chief of npj Systems Biology and Applications. He was not involved in the journal’s review of, or decisions related to, this manuscript.

Ravi Salgia and Prakash Kulkarni are co-guest editors of the series titled “Phenotypic Plasticity: A Systems Approach” npj Systems Biology and Applications. They were not involved in the journal’s review of, or decisions related to, this manuscript.

---

**Fig. S1.**
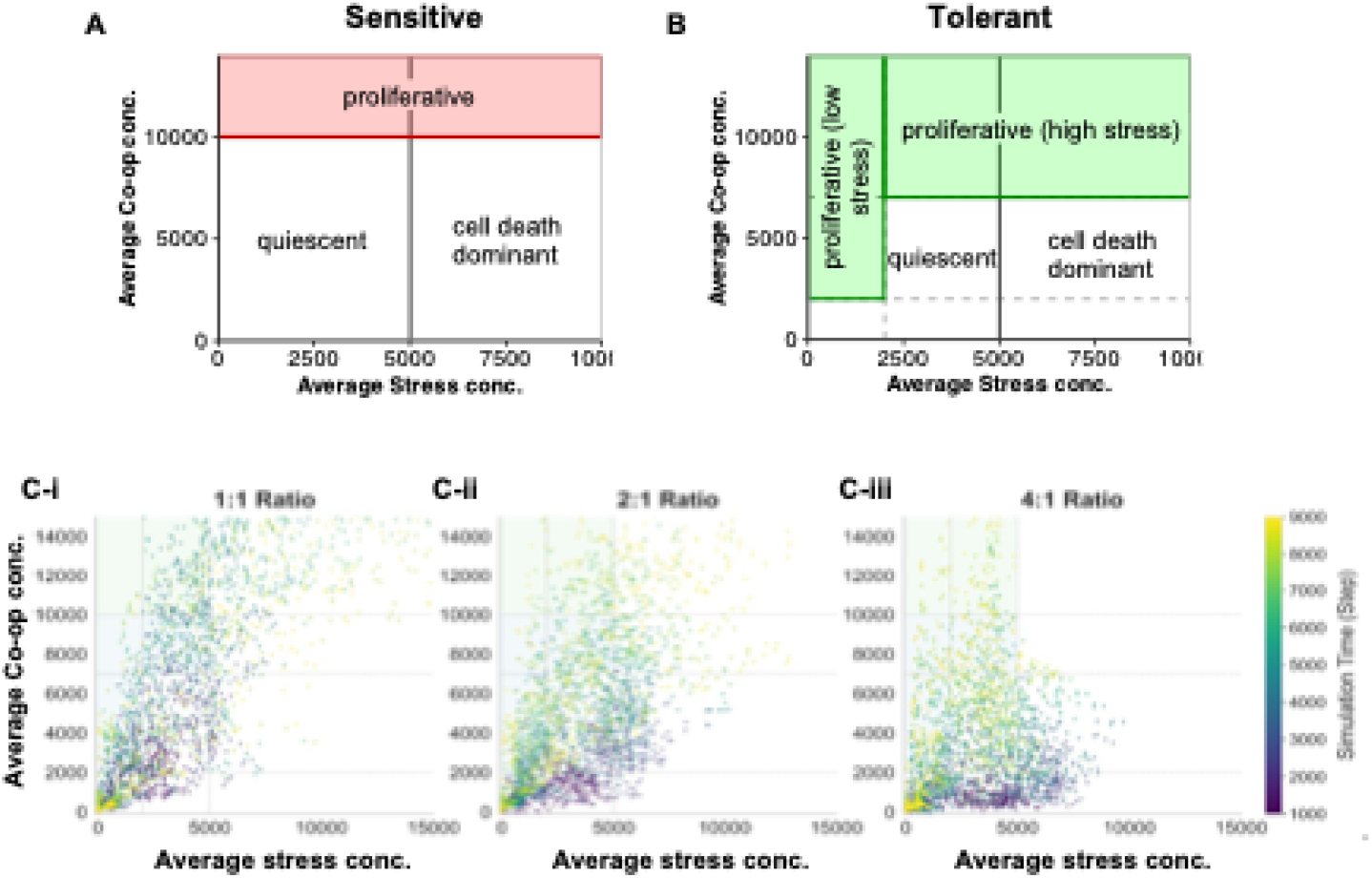
(A-B) Schematics showing different ecological regimes encountered by the sensitive and tolerant cells during simulation. The zone labels are based on cell behavior determined by predefined coop and stress thresholds; proliferative zones where cell division supersedes cell death, leading to net proliferation; quiescent: no division or death; (C) Scatter plots showing the covariation of average stress and co-op concentrations, with data points colored according to simulation timepoint (MCS). Each panel represents simulations at a given seeding ratio.

**Fig. S2.**
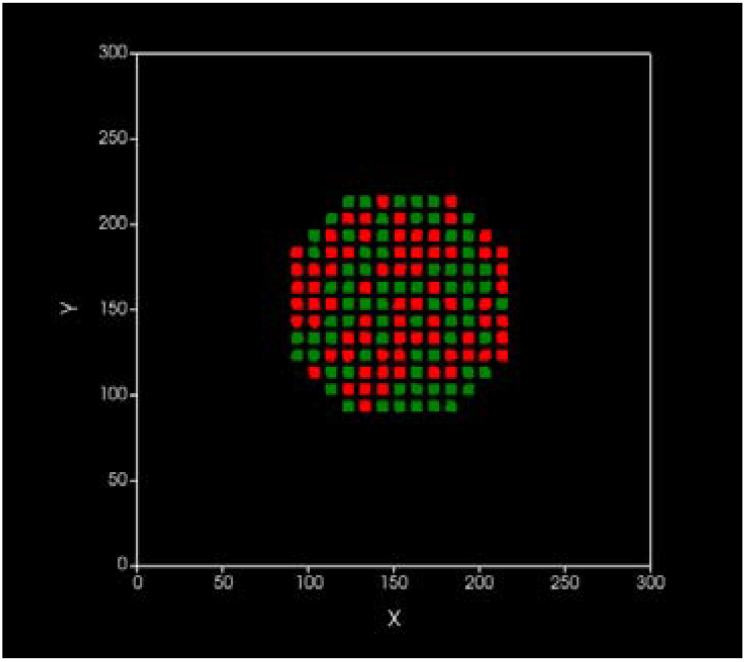
Example starting configuration for multiscale simulations showing sensitive and tolerant populations in red and green respectively.

